# ASLPrep: A Generalizable Platform for Processing of Arterial Spin Labeled MRI and Quantification of Regional Brain Perfusion

**DOI:** 10.1101/2021.05.20.444998

**Authors:** Azeez Adebimpe, Maxwell Bertolero, Sudipto Dolui, Matthew Cieslak, Kristin Murtha, Erica B. Baller, Bradley Boeve, Adam Boxer, Ellyn R. Butler, Phil Cook, Stan Colcombe, Sydney Covitz, Christos Davatzikos, Diego G. Davila, Mark A. Elliott, Matthew W. Flounders, Alexandre R. Franco, Raquel E. Gur, Ruben C. Gur, Basma Jaber, Corey McMillian, the ALLFTD Consortium, Michael Milham, Henk J.M.M. Mutsaerts, Desmond J. Oathe, Christopher A. Olm, Jeffrey S. Phillips, Will Tackett, David R. Roalf, Howard Rosen, Tinashe M. Tapera, M. Dylan Tisdall, Oscar Esteban, Russell A. Poldrack, John A. Detre, Theodore D. Satterthwaite

## Abstract

Arterial spin labeled (ASL) magnetic resonance imaging (MRI) is the primary method for non-invasively measuring regional brain perfusion in humans. We introduce ASLPrep, a suite of software pipelines that ensure the reproducible and generalizable processing of ASL MRI data.

## MAIN

Despite comprising only 2% of the human body mass, the adult brain receives approximately 15% of cardiac output to support the intensive demands of neural computation^1^. Cerebral blood flow (CBF) is tightly linked to brain metabolism^2^, varies predictably across the lifespan^3^, and is increasingly seen as an important biomarker of diverse neuropsychiatric and neurological illnesses^4^. Although the gold standard method of measuring CBF is 15O-PET, arterial spin labeled (ASL) perfusion magnetic resonance imaging (MRI) has evolved to become the dominant method for non-invasive measurement of CBF in humans due to its ease of implementation, lower cost, and lack of ionizing radiation^5^.

The ascendancy of ASL MRI has also been accompanied by a rapid rise of both acquisition methods and analytic techniques^5^. For example, widely used ASL MRI sequences vary in their labeling type, number of echo times, labeling duration, number of post-labeling delays used, image scaling, background suppression, and whether or not a reference (*M0*) image is acquired. Furthermore, different MRI schemes for ASL may yield markedly different output: commonly used schemes can provide a timeseries of control and label pairs, a single difference image, or a fully quantified CBF image. When combined, these factors have limited the generalizability of techniques for the processing and quantification of CBF and have slowed the pace of translational research^6^. To address this gap, we introduce ASLPrep: a generalizable and robust software workflow that allows for reproducible processing of a wide range of ASL MRI data (**Figure 1**).

**Figure 1.**
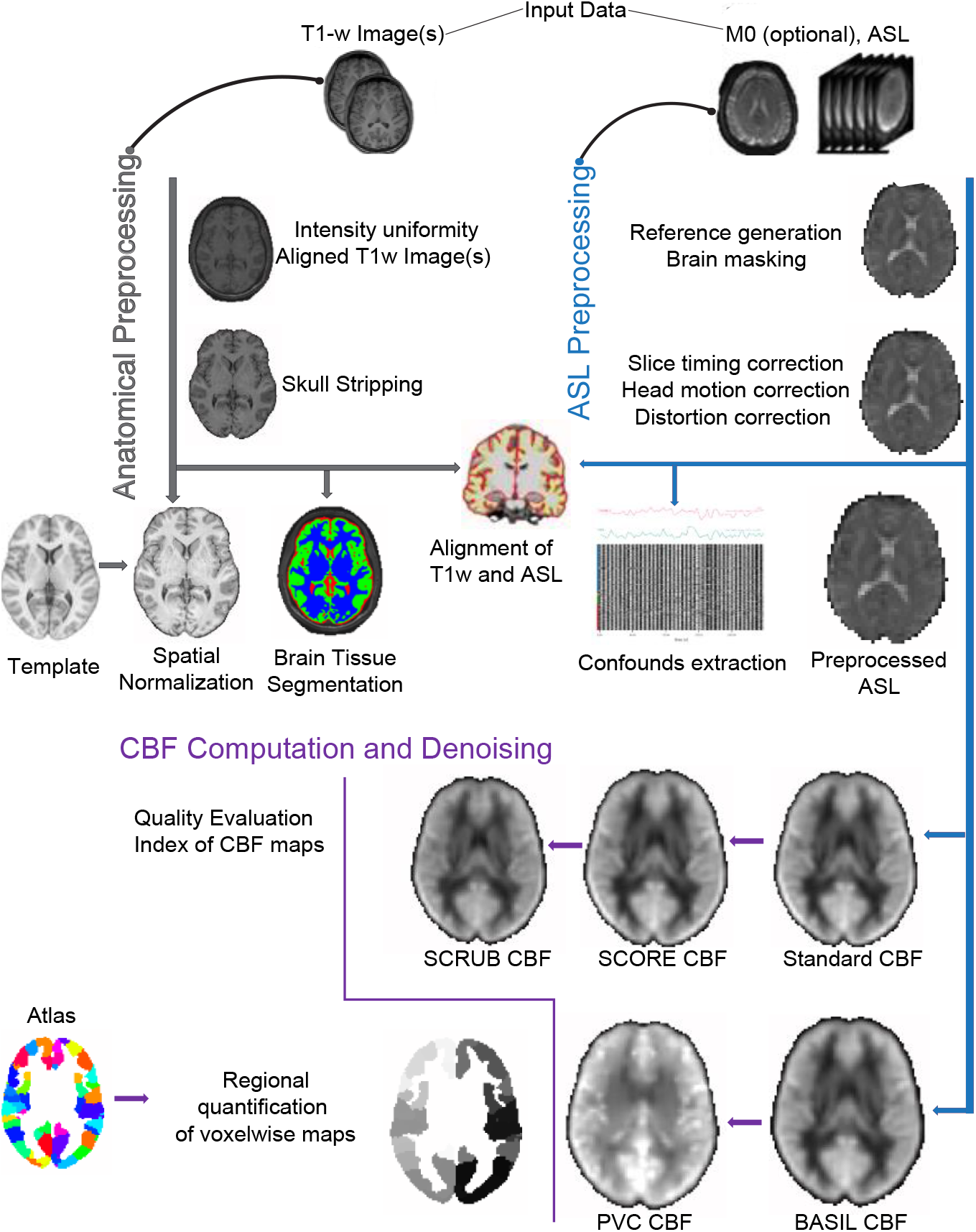
Overview of ASLPrep. Input data to ASLPrep include ASL images, anatomical (T1 weighted) images, and (optionally) M0 reference images. Anatomical preprocessing is executed using standard tools (as implemented in sMRIPrep), while ASL preprocessing is done with ASLPrep. CBF computation and denoising can be executed using multiple options, including the standard CBF procedure, and can optionally include SCORE, SCRUB, and BASIL with or without partial volume correction (PVC). Importantly, ASLPrep generates extensive quality indices, as well as a visual report of each step. Finally, regional CBF is summarized according to standard or custom atlases.

ASLPrep requires the metadata be recorded in Brain Imaging Directory Structure (BIDS)^7^ format and leverages BIDS to automatically configure appropriate workflows based on the data provided. ASLPrep is designed with a focus on reliability, and builds upon widely-used neuroimaging toolboxes, such as FSL^8^, FreeSurfer^9^, AFNI^10^, and ANTs^11^ (**Supplementary Table 1**). ASLPrep also includes in-house implementations for algorithms unavailable elsewhere, for instance the SCORE de-noising option^12^, which is particularly useful for studies of populations with greater head motion, such as children and many patient groups.

Building on this preprocessing workflow, ASLPrep can optionally execute advanced methods to quantify CBF. In addition to the standard CBF quantification procedure, ASLPrep includes two different Bayesian models that incorporate information regarding brain structure: (BASIL^13^ and SCRUB^14^). ASLPrep also allows users to specify if they would like to implement partial volume correction^15^ (with BASIL), which adjusts CBF according to the mixture of gray and white matter present in the anatomical image. For all quantification models, regional CBF is summarized in a diverse set of bundled atlases or custom atlases provided by the user.

Both minimal preprocessing and quantification workflows are transparently documented with a detailed visual report that generated dynamically. Critically, each step in the workflow is demonstrated and its performance can be assessed for quality with “before vs. after” visualizations (see **Supplementary Figure 1**). In addition to such visualizations, ASLPrep provides multiple quantitative measures of image quality (see Supplement). Like fMRIPrep^16^, these visual reports also include a “citation boilerplate” that comprehensively describes the actual workflow implemented, including software versions and relevant citations to facilitate maximally transparent reporting in papers that use ASLPrep.

ASLPrep is distributed as a Docker image that includes all dependencies (https://hub.docker.com/r/pennlinc/aslprep), ensuring that it can be run in nearly any computing environment. The modular code base of ASLPrep uses Nipype^17^ and is openly developed on GitHub (https://github.com/pennlinc/aslprep), allowing for rapid detection of bugs, integration of feature requests, and support for the international user base. Prior to the release of patches or new versions, all changes to the underlying code of ASLPrep are subject to continuous integration testing via CircleCI. Extensive documentation (https://aslprep.readthedocs.io) is version-controlled and frequently updated, facilitating broad dissemination. A total of more than 17,000 data have been successfully run through ASLPrep.

To illustrate the generalizability of ASLPrep, we processed five different datasets acquired with a wide range of acquisition parameters (n=3,150 total scans; see **Supplementary Table S2**). These datasets included four ASL sequences collected on Siemens scanners using pseudo-continuous labeling (e.g. pCASL), but different encoding schemes: 2D spin echo PCASL images from the Philadelphia Neurodevelopmental Cohort (PNC; n=1,491), 2D gradient echo images from the NKI-Rockland Sample (NKI; n=1,257), a 3D stack-of-spirals spin echo acquisition from a study of irritability in youth (IRR; n=115), and a publicly available study of aging that used a 2D spiral gradient echo PCASL sequence (AGE; n=63). Furthermore, we included a study of fronto-temporal dementia (FTD; n=110) that was collected on a GE scanner using a 3D EPI gradient echo sequence with pCASL. Of these studies, two included a reference (M0 scan; IRR and FTD), and one (FTD) provided a difference image (ΔM) rather than full label/control timeseries. For each of these diverse datasets, we completed both minimal preprocessing and CBF quantification. Specifically, we evaluated the mean CBF of gray matter and white matter in each dataset, and how gray matter CBF evolved with age^18^. While these analyses focused on CBF quantified using the standard CBF quantification procedure, supplementary analyses detail results from the other quantification procedures packaged with ASLPrep.

Across both datasets and workflows, pipelines automatically configured by ASLPrep concluded without run time errors. As part of quality assurance, 4% participants with gross motion (frame-wise displacement [FD] greater than 1mm) or non-physiologic CBF (e.g., a ratio of GM:WM CBF of less than 1) were excluded from further analyses (see **Supplementary Table 3**; final sample n=3,021). Inspection of data from individual participants (see exemplar **Supplementary Figure 2**) as well as group average CBF from each dataset (**Supplementary Figure 3**) revealed consistent performance. As expected, the distinction between gray matter and white matter CBF was more striking in datasets of youth and was reduced in datasets that were composed of older individuals (**Figure 2A** and **Supplementary Figure 4**). When data from individual participants were aggregated across all datasets, the anticipated nonlinear decline of CBF over the lifespan was clearly evident (**Figure 2B**; also see **Supplementary Figure 5**).

**Figure 2.**
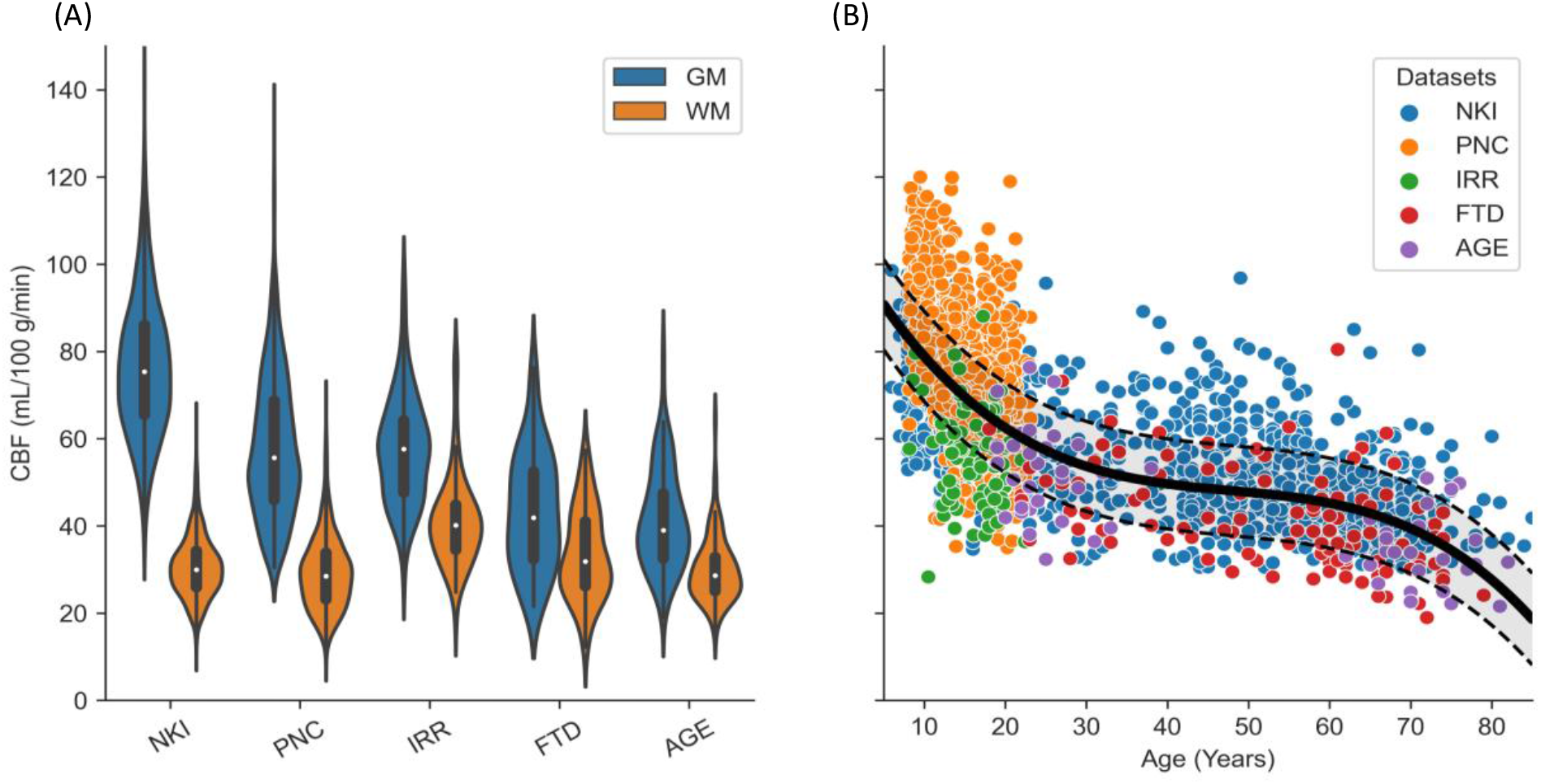
ASLPrep provides generalizable pre-processing across sequences, scanners, and the lifespan. (**A)** Cerebral blood flow (CBF) within gray matter (GM) and white matter (WM) for each dataset. CBF is higher in GM than in WM in all datasets, but this difference is attenuated in studies of older adults, including those with fronto-temporal dementia (FTD). (**B**) GM CBF declines predictably across the lifespan. Data aggregated across all datasets reveals a predictable pattern when modeled using generalized additive models with penalized splines. GM CBF declines steeply in youth, is relatively stable during adulthood, and declines again after age 60.

Across the over 3,000 participants evaluated, the processing time of ASL images never exceeded 40 minutes (when executed using 4 cores and 30 GB RAM) Nonetheless, the total runtime of ASLPrep was substantially longer (mean 2.5 hours), driven largely by its dependencies on anatomical image processing^19^ (see **Supplementary Figure 6**). However, one critical feature of ASLPrep is that it can consume processed anatomical images that conform to the BIDS-derivatives standard (e.g., sMRIPrep^19^) obviating the need for re-processing structural images and dramatically accelerating runtime. This key feature is particularly important for multi-modal imaging studies as it allows a unique source of preprocessed anatomical information can be used, ensuring consistency across image types (ASL, fMRI, dMRI, etc).

Two limitations of the current version of ASLPrep should be noted. First, although ASLPrep provides a comprehensive framework for evaluating perfusion quantification workflows, such an evaluation is beyond the scope of this paper. Second, certain experimental ASL acquisition schemes (such as QUASAR^20^) include parameters that are not currently supported by BIDS; as such, these schemes cannot currently be processed by ASLPrep.

In summary, ASLPrep allows investigators to correctly apply reproducible preprocessing pipelines and advanced CBF quantification methods to nearly all ASL images. ASLPrep adapts its workflow to the characteristics of the input data, ensuring appropriate image processing as long as the data has been correctly specified in BIDS. By harnessing complementary techniques from multiple software packages and combining them in an interoperable framework, ASL reduces the burden on investigators who wish to avoid learning the details of many disparate techniques. Taken together, ASLPrep ensures fully reproducible and widely generalizable processing, quality assurance, and quantification of ASL images.

## Supporting information

https://upenn.box.com/s/vziw20savmzffewvla7qny5zltw7667f

## SOFTWARE AVAILABILITY

ASLPrep is available under the BSD 3-clause license at https://github.com/pennlinc/aslprep. Docker images corresponding to every new release of ASLPrep are automatically generated and made available on Docker Hub. All code used to perform the statistical tests are available at: https://pennlinc.github.io/aslprep_paper, under the BSD-3-Clause License.

## DATA AVAILABILITY

Imaging data are available with restrictions depending on the original source of the data. PNC data are available on dbGAP [https://www.ncbi.nlm.nih.gov/projects/gap/cgi-bin/study.cgi?study_id=phs000607.v3.p2]. NKI neuroimaging data are openly available on the NeuroImaging Tools and Resources Collaboratory [http://fcon_1000.projects.nitrc.org/indi/enhanced/]. IRR and FTD data are available upon request to TDS and CM respectively. AGE data are available on Open Neuro [https://openneuro.org/datasets/ds000240/versions/00002].

## ETHICS OVERSIGHT

No new data were collected specifically for this study. All other data were acquired with IRB approval at their original institutions. The University of Pennsylvania IRB approved the PNC, FTD, and IRR studies. The AGE and NKI datasets are publicly available, de-identified data resources.

