## Supplementary material for "ASLPrep: A Generalizable Platform for Processing of Arterial Spin Labeled MRI and Quantification of Regional Brain Perfusion": https://upenn.box.com/s/vziw20savmzffewvla7qny5zltw7667f

#### ONLINE METHODS

ASLPrep allows investigators to easily process diverse arterial spin labeled (ASL) MRI[1,2] data and compute cerebral blood flow (CBF). ASLPrep is designed using an adaptive architecture that leverages the Brain Imaging Data Structure (BIDS)[3], an open standard for describing neuroimaging data. ASLPrep reads the meta-data provided by BIDS, allowing workflows to automatically adapt to the parameters of the data without manual intervention. ASLPrep leverages Nipype[4] to ensure compatibility across tools from many software packages (e.g., Freesurfer, AFNI, ANTS, FSL) and effectively combine their complementary strengths (see **Supplementary Table 1**). ASLPrep's design is inspired by fMRIPrep[5] and adheres to the principles of NiPreps[6] (www.nipreps.org): it maximizes interoperability and adaptability to input data with BIDS, it reproducibly delivers "analysis-ready" data so that researchers can confidently focus on statistical modeling, and code is managed with software engineering techniques to ensure quality and reliability, following BIDS-Apps' directions[7] (e.g., utilizes open-source development, implements version control with GitHub, and includes continuous integration of testing with CircleCI to check every code update, etc.). ASLPrep is composed of four main workflows (**Figure 1**): anatomical preprocessing, ASL preprocessing, CBF computation, and assessment of image quality.

##### Anatomical Preprocessing

The anatomical preprocessing workflow in ASLPrep leverages sMRIPrep (version 0.6.1)[8], a structural magnetic resonance imaging (sMRI) processing pipeline. sMRIPrep performs basic processing steps including subject-wise averaging, bias field correction, segmentation, and spatial normalization. The anatomical outputs of sMRIPrep can be used while processing multi-modal imaging data, including fMRIPrep[5], QSIPrep[9], and ASLPrep.

The major steps of anatomical preprocessing are summarized below:

*Bias field correction and T1w subject-wise averaging*: The structural T1w image is first corrected for intensity nonuniformity with *N4BiasFieldCorrrection*[10] as implemented in ANTs. If there are several T1w images, bias-corrected corrected images are fused into a reference T1w map with Freesurfer's *mri_robust_template*[11].

*Brain extraction and tissue segmentation*: The bias-corrected T1w image is skull-stripped with *antsBrainExtraction.sh* using either the OASIS[12] (default) or NKI[13] template. Note that the template used here for brain extraction is distinct from the template used for spatial normalization. After brain extraction, FSL's *FAST*[14] is used to segment the T1w brain into cerebrospinal fluid (CSF), grey matter (GM), and white matter (WM). FAST produces both a hard segmentation as well as partial volume estimates for each tissue class.

*Spatial normalization and template selection:* Following bias correction and brain extraction, the T1w image is normalized to the MNI152 Nonlinear Asymmetric[15,16] template using the top-performing deformation provided by *antsRegistration*[17]. Beyond the MNI template, Templateflow[16] allows for the selection of other available templates including the PNC[18], NKI, and OASIS templates.

**ASL Preprocessing**

Due to the variety of ASL data types, ASL preprocessing workflows require that the input data conform to the ASL BIDS specification[7]. The processing workflows can accommodate disparate ASL labeling approaches[1],readout methods[19], and number of volumes in the ASL input data. The most common labeling approaches include continuous labeling (CASL), pulsed labeling (PASL), and pseudo-continuous labeling (pCASL). CASL was the first popular ASL labeling technique; however, one of its major drawbacks is that it requires long radio frequency (RF) labeling pulses, which causes a magnetic transfer (MT) effect that confounds the ASL signal. PASL uses short RF pulses and is thus less susceptible to MT effects, but suffers from low sensitivity compared to CASL. The pCASL method was developed as an intermediate technique that takes advantage of CASL's high signal to noise ratio (SNR) and PASL's high labeling efficiency, without the need for long labeling pulses. pCASL achieves this using a train of short RF pulses rather than continuous RF[20].

*Input data*: The four possible types of ASL data provided by an acquisition sequence are (a) a timeseries of control and label images, (b) a $\Delta M$ image, (c) a *M0* image, and (d) a quantified CBF image. It should be noted that the $\Delta M$ and CBF images are derived rather than raw data. However, some ASL sequences from GE or Philips scanners provide these derived images instead of the raw ASL timeseries; as such, -they are considered as potential input data to ASLPrep. These possible inputs are briefly defined below:

A. ASL timeseries. A typical ASL timeseries consists of control and label images, which are acquired in pairs in an ASL sequence, and scaled identically. When background suppression is not used, the control image can be used in place of the *M0* image for calibration. Most but not all ASL sequences will provide this timeseries; some sequences will provide an $\Delta M$ image or the fully-quantified CBF image.
B. $\Delta M$: The $\Delta M$ images are formed by the pairwise subtraction of the label and control images. Multiple control-label pairs are always acquired, which after subtraction generates a timeseries of $\Delta M$ images. Some sequences provide an average $\Delta M$ image rather than the full ASL timeseries of control and label pairs; ASLPrep can recognize and process either.
C. *M0*: A *M0* image is used as a reference image and to estimate the equilibrium magnetization (M0) of blood. If a *M0* scan is not provided, the average of the control images is used as

the reference (while checking for background suppression). Alternatively, a user-specified *M0* value specified in the BIDS meta-data will override the *M0* scan (if present) and be used for calibration.

D. CBF: The quantified cerebral blood flow (CBF) image is produced by dividing the $\Delta M$ by the processed *M0* image or by the user specified *M0* value specified in BIDS. Using a standard model, the CBF image is then scaled into physiological units (mL/100g/min). Rarely, ASL sequences provide a fully quantified CBF map instead of an ASL timeseries or a $\Delta M$ image. In this case, most steps of ASLPrep are obviated. However, normalization to a template and calculation of average values for atlas parcels can still be performed. In this case, an *M0* image is necessary for generation of a high-quality brain mask.

Using this input data, the general ASL preprocessing workflow is as follows:

*Reference volume selection*: For each ASL dataset, a reference volume for motion correction and co-registration is selected. When an ASL timeseries is provided, the median of ASL volumes is selected as the reference volume. For a single $\Delta M$ or CBF image, the *M0* scan (if present) is used as the reference volume. The reference volume is skull-stripped with FSL's *BET*[21] and refined with the co-registered T1w brain mask.

*Motion estimation*: The head motion parameters from the ASL timeseries (if available) are estimated with FSL's *MCFLIRT*[22].

*Slice timing correction*: If slice-timing information is available in the metadata, the slice-timing correction is performed on the ASL data with AFNI's[23] *3dTshift* . However, slice-timing correction is optional, can be turned off, and is not applied if slice times are not specified in the BIDS meta-data. Note that while slice timing correction can be applied for 2D ASL data, it is not recommended for 3D ASL data.

*ASL-T1w Co-registration*: Co-registration aligns the T1w image and ASL reference volume. This uses boundary-based co-registration as implemented with FSL's *FLIRT*[24]. A rigid body transform (6 DOF) is specified by default. Forward and backward transformation matrices are generated for required subsequent preprocessing.

*Distortion correction*: Distortion correction is implemented using SDCFlows[25] (Susceptibility Distortion Correction workFlows). SDCFlows provides workflows for the preprocessing of several MRI schemes that allow the estimation of $B_0$ field-inhomogeneity maps, which are directly related to the distortion. Distortion correction is applied to the ASL data if the appropriate fieldmap information is provided in the BIDS meta-data. Distortion correction is optional. SDCFlows additionally includes an experimental fieldmap-less[26] distortion correction method, which uses a nonlinear registration between the ASL reference image and the T1w image.

#### CBF Computation

Following common preprocessing steps, there are two streams for CBF computation and denoising. The first stream is the default (Basic CBF) and consists of CBF computation with the general kinetic model[27,28]. The first step produces a CBF timeseries if there are multiple control-label pairs or $\Delta M$ images provided. Optionally, this CBF timeseries can then be de-noised using Structural Correlation based Outlier REjection (SCORE)[29]. Last and also optionally, a Bayesian model can be applied that incorporates data from the anatomic image using Structural Correlation with RobUst Bayesian (SCRUB)[30].

The second stream uses FSL's Bayesian Inference for Arterial Spin Labeling (BASIL) toolbox. BASIL's Bayesian model incorporates spatial regularization and produces a single CBF map. (Note that because it produces a single CBF map rather than a CBF timeseries, it cannot be used jointly with SCORE). Following computation of the CBF map, BASIL can also optionally calculate a CBF map with partial volume correction (PVC) applied.

Both common steps and specific procedures for each of the two CBF computation streams are detailed below:

<u>Procedures common to both processing streams</u>

*$\Delta M$ Computation*: After ASL preprocessing, the $\Delta M$ images are extracted. Depending on the ASL input data, the label and controls images are subtracted pairwise to obtain a timeseries of $\Delta M$ images. However, if input data contains a $\Delta M$ image or CBF maps, this step is not performed.

*CBF calibration:* As noted above, the $\Delta M$ images require scaling in order to calculate CBF. This can be done using either a dedicated *M0* image (preferred), the average of the control images from the ASL timeseries (if no background suppression is used), or a user-specified value provided in the JSON. If an *M0* image is present, it will be used instead of the mean of the control images. However, specification of an *M0* value in the JSON will force its application regardless of the other images present; this should generally be avoided. Default smoothing of M0 scan images is implemented with 5mm full width at half maximum (FWHM), but the users can adjust to any value. Smoothing of the *M0* is advisable for CBF computation.

<u>Stream 1: Standard CBF +/- SCORE and SCRUB</u>

*CBF quantification with the standard model*: CBF is quantified based on the general kinetic model[27,28] and requires several parameters[1], including labeling duration (LD), post labeling delay (PLD), T1-blood relaxation time, labelling efficiency, blood-brain partition coefficient, and inversion time. Some of these parameters (like bolus duration, LD, and PLD) are specific to the ASL labelling techniques. The other parameters (like labelling efficiency and blood-brain partition coefficient) depend on the labelling approaches (PCASL or PASL) and hardware (e.g., MRI machine magnetic strength). ASLPrep quantifies CBF for both single delay (one PLD) and multi-delay (multiple PLDs) ASL data. The multi-delay ASL data gives us the opportunity to estimate arterial transit time (ATT) using signal weighted delay[31]. All parameters necessary for computation of CBF using the standard model are read from the JSON that adheres to the BIDS specification for ASL.

*CBF Denoising with SCORE*: CBF is known to be susceptible to artifacts[19,32] especially from head motion. ASL's low signal to noise ratio (SNR), particularly when acquired without background suppression, is compensated for by averaging multiple label-control pairs. However, some volumes may be corrupted and can influence the average of the CBF map. The corrupted volumes are likely to be outliers of the CBF timeseries. To identify and remove these outlier volumes, ASLPrep implements Structural Correlation based Outlier REjection (SCORE)[29]. The first step of SCORE is the detection of a small number of extreme outliers before further processing. These outliers are identified as CBF volumes with mean CBF within grey matter (GM) tissue that is greater than 2.5 times the standard deviation of the mean GM CBF. In the second step, each remaining CBF volume is compared to average CBF maps in order to detect noise in these volumes. The variability of CBF values within the three tissue types, GM, WM and CSF is calculated. Then, CBF volumes with high variance of CBF values within the three anatomical tissues are flagged as outliers. In each step, outliers are identified, and the remaining CBF volumes are averaged. The variance of averaged CBF within the three anatomical tissue classes is then compared with the previous iteration. The procedure is repeated while the CBF variance within anatomical tissues is lower than that previous iterations. In the last iteration, all outlier volumes are discarded before averaging the remaining CBF volumes. For further details see Dolui et al 2017[29].

*Bayesian Estimation of CBF with SCRUB*: While SCORE detects and removes outlier volumes that contribute to spatially constrained artifacts, Bayesian techniques may be used to improve SNR. Accordingly, ASLPrep optionally allows users to implement Structural Correlation with RobUst Bayesian (SCRUB)[30]. SCRUB incorporates information from the structural image as a prior in a Bayesian to reduce noise and improve SNR. SCRUB uses an iterative reweighted least square method to estimate CBF within a Bayesian framework. In this framework, the reweighted CBF at each voxel is compared to the prior provided by a standard CBF based on the tissue probability maps from the soft segmentation of the anatomic image. A reliable CBF map is more likely when the CBF temporal variance is less than the mean CBF variance. If the CBF temporal variance is greater than the mean CBF variance, a higher weight is assigned to the prior term. For further details regarding SCRUB, see Dolui et al 2016[30].

Stream 2: CBF Computation with BASIL +/- Partial Volume Correction

*CBF Computation with BASIL:* ASLPrep includes the ability to compute CBF using BASIL[33] (Bayesian Inference for Arterial Spin Labeling). BASIL was originally developed for CBF computation of multi-PLDs data, but it also can be used with single PLD data. BASIL uses a fast Bayesian inference method for the kinetic model inversion and includes perfusion estimation and associated variables, such as arterial transit time. BASIL also allows for the inclusion of the variability of other model parameters, such as relaxation times for tissue and blood, as well as labelling durations. After CBF computation, BASIL applies a gaussian process based prior to computed CBF maps for adaptive spatial regularization[34]. For further details regarding BASIL, see Chappell et al 2009[33].

*Partial Volume Correction with BASIL:* The low spatial resolution is a limitation of ASL and leads to partial volume effects[1]. Partial volume effects occur when voxels near the boundary between different tissue types (e.g., GM and WM) contain a mixture of the respective tissues. As a result, a given voxel may have apparently lower CBF due to the greater proportion of WM at that location. Partial volume effects are particularly relevant in ASL data, where the ASL signal intensity at each voxel represents mixtures of signals from GM and WM (CSF perfusion is assumed to be zero). Partial volume effects can be accounted for using high resolution GM and WM probability maps. The high-resolution GM and WM probability maps are transformed into the low-resolution space of the ASL data, allowing the relative mix of GM and WM at each ASL voxel to be estimated. BASIL uses this data in conjunction with adaptive spatial regularization to yield partial volume corrected (PVC) CBF maps. For further details, see Chappell et al 2011[35].

**Measures for Quality Control**

ASLPrep generates a rich set of indices to assist in the quality control process. Because many errors of image processing result from problems with image registration, ASLPrep calculates measures that assess the quality of both co-registration of the ASL to the T1w image, as well as normalization of the T1w image to the specified template. Specifically, ASLPrep calculates the mask overlap, spatial correlation, Dice coefficient, and Jaccard index for each step of registration. Furthermore, ASLPrep provides several quality measures for the ASL timeseries, including the mean framewise displacement (FD)[36], the root mean square variance of temporal derivative of CBF time courses (DVARS)[37], the number of voxels with a negative CBF, and the CBF Quality Evaluation Index (QEI)[38]. QEI is a quantitative metric of the quality of CBF maps based on the CBF map's similarity with the structural tissues, the CBF spatial variability within each tissue class (GM, WM), and the percentage of negative voxels within the GM mask. QEI ranges from 0 from 1, with higher values referring to higher quality CBF maps; prior work has established that it is an excellent proxy of CBF image quality[38]. Finally, ASLPrep calculates the ratio of CBF within the GM mask to CBF within the WM mask; this ratio is expected to be greater than 1. All quality indices described above are written to a TSV for each session processed; these are easily concatenated across subjects to facilitate rapid quality assurance of large-scale ASL studies.

**Regional Quantification**

As a final step, ASLPrep optionally quantifies the mean CBF within each parcel within standard atlases. To do this, standard atlases are transformed to CBF native space using a single interpolation. The mean of CBF values of each atlas's parcels is extracted into a comma-separated values (csv) file. At present, the Harvard-Oxford[39] and Schaefer[40] (200 and 400 parcel resolution) atlases[39] are included.

**Standard Output**

Processed data are named and include meta-data to conform to the according to the proposed BIDS specification for derived data.

*Anatomical derivatives:* Anatomical derivatives generated by sMRIPrep are placed in each subject's *anat/* subfolder. The major derivatives, in T1w and template spaces, include:

a. Preprocessed T1w, brain mask, and tissues segmentation mask
b. GM, WM, and CSF partial volume estimates
c. Transformation files for normalization between the T1w image and the template

*ASL derivatives:* Similarly*,* ASL derivatives are stored in the *perf/* subfolder. Based on the anatomical preprocessing, users can specify one or more output spaces: native ASL, T1w, and one or more standard templates (MNI, OASIS, PNC and others, as available on *TemplateFlow*[16]*)* MNI output space is the default, which is written out in the resolution of the original ASL image. However, users can specify as many as output spaces as they want. The major output in any space includes:

- ASL reference volume, brain mask, and preprocessed data
- CBF timeseries and mean CBF map
- Transformation files or transforms between T1w and ASL reference
- Arterial transit time image *if multiple-PLDs are available*
- SCORE and/or SCRUB CBF maps *if requested*
- BASIL and/or PVC CBF maps *if requested*
- Quality control measures summarized in TSV file. The quality control measures include mean frame-wise displacement (FD), relative root mean square of motion parameters (relRMS), DVARS, registration and normalization indexes, mean CBF, ratio of $CBF_{GM}$ to $CBF_{WM}$ and quality evaluation index for each CBF map.
- Regional quantification of CBF according to specified atlases, written in a CSV file.

- Confound matrix for each ASL volume in TSV file. The confound matrix includes six motion parameters, FD, and DVARS for each processed ASL volume.

**ASLPrep Report**

ASLPrep generates a descriptive HTML report for each subject and session (see **Supplementary Figure 1**). The report begins with a summary of key parameters found by ASLPrep in the BIDS layout. Subsequently, it lists the key operations and processing steps applied to the dataset. Notably, each step includes a thorough visual assessment of the data, including "before" versus "after" animations of each step. These visualizations include normalization, co-registration, and distortion correction. The report additionally includes a carpet plot of both the raw and processed image timeseries, as well as views of all CBF maps generated. Importantly, the report details multiple quality control features, including in-scanner motion, QEI, co-registration and normalization quality (overlap of coverage, Dice coefficient, Jaccard index), and mean CBF within both gray and white matter masks. Critically, the report ends with boilerplate methods text, which provide a clear and consistent description of all preprocessing steps used, provided with appropriate citations.

**Application of ASLPrep to Lifespan Data**

*Datasets and ASLPrep Execution*

The general workflow (see **Figure 1**) was applied to datasets with diverse subject populations and scanning parameters. **Supplementary Table 2** describes the acquisition parameters of the datasets used; **Supplementary Table 3** details the major preprocessing operations that were automatically applied to each dataset. One dataset included both an ASL timeseries of control-label pairs as well as an *M0* image (IRR). In contrast, three datasets had control-label timeseries but lacked an *M0* image (NKI, PNC, AGE), and one dataset only included $\Delta M$ and *M0* images (FTD). Furthermore, the acquisition types of these datasets were also different: PNC and NKI were acquired in 2D while the other datasets were acquired in 3D. All data were collected using 3T scanners. For each study, CBF was quantified using four approaches: the standard CBF model, SCRUB (with SCORE denoising turned on), BASIL, and BASIL + PVC. The only exception to this was the FTD dataset, where SCRUB could not be applied as it requires an ASL timeseries.

Each subject data was processed using ASLPrep version 0.2.6[41] with 4 cores and 30 GB RAM on the CUBIC High-Performance Cluster (HPC) at the University of Pennsylvania. Notably, the anatomical preprocessing with sMRIPrep required substantially more time than ASL processing (**Supplementary Figure 6**). While anatomical preprocessing took an average of 4.4 hours, the ASL processing was completed in an average of 30.78 minutes.

*Statistical Analysis*

All ASL data outputs were resampled to MNI2009a with resolution of 2mm for uniformity. For each CBF map, the mean CBF within GM and WM masks were extracted. As part of quality assurance, we excluded participants with mean FD greater than 1 mm or a CBF GM:WM ratio of less than 1. The quality assurance process resulted in a sample of 3,138 participants used for subsequent analysis (see **Supplementary Table 3** for more information). CBF within GM and WM masks were extracted for all the CBF maps (standard CBF, SCRUB, BASIL, and BASIL+PVC). In order to rigorously model both linear and nonlinear evolution of CBF over the lifespan, the mean GM CBF was regressed on age using a generalized additive model (GAM) with penalized splines.

**SUPPLEMENTARY TABLES**

**Supplementary Table 1: Software used by ASLPrep.** ASLPrep integrates diverse software in its processing workflow.

| Processing step | Implementation | Notes |
|---|---|---|
| *Input data* | | |
| Data in NIfTI format | BIDS format | anat, asl, fmap(optional) |
| *Anatomical processing (with sMRIPrep)* | | |
| T1w alignment | | If more than T1w images |
| Bias correction | ants's N4 | Required |
| Skull stripping | antsBrainExtraction | Required |
| Segmentation | FSL's FAST | Required |
| Spatial Normalization | antsRegistration | Required |
| *ASL processing* | | |
| Reference generation | new implementation | Required |
| Slice timing correction | AFNI's 3dTShift | Optional |
| Head motion estimation and correction | FSL's MCFLIRT | Required |
| Distortion correction | SDCFlow | Optional |
| Registration | FSL's FLIRT with BBR | Alignment of T1w and ASL reference |

| | | |
|---|---|---|
| Confounds Extractions | Niworkflows[41] | six motion parameters and dvars |
| *CBF computation* | | |
| M0 processing | new implementation | Masking and smoothing.<br>Required |
| CBF quantification | new implementation | Required |
| BASIL | FSL | Bayesian Inference for Arterial Spin Labeling (BASIL):<br>Alternative to CBF computation with spatial regularization<br>Optional |
| *CBF post processing (optional)* | | |
| SCORE | new implementation | Structural Correlation based Outlier Rejection (SCORE):<br>discard few extreme outliers |
| SCRUB | new implementation | Structural Correlation with Robust Bayesian (SCRUB):<br>Voxel-wise empirical robust Bayesian estimation |
| BASIL PVC | FSL | Partial volume correction (PVC) |
| *Other processing* | | |
| Quality control | new implementation | Motion summary measures (FD and relRMS)<br>Registration and Normalization quality indices<br>Quality evaluation index and Mean CBF (GM, WM, and ratio) |
| Regional quantification | new implementation | Extract CBF with atlases |
| Visual Report | Modified Niworkflows | Detail processing reports and boilerplate generation |

#### Supplementary Table 2: Evaluation data

| Parameter/Data | IRR | PNC | NKI | AGE | FTD |
|---|---|---|---|---|---|
| **Manufacturer** | Siemens | Siemens | Siemens | Siemens | GE |
| **MR acquisition type** | 3D | 2D | 2D | 3D | 3D |
| **M0scan** | Yes | No | No | No | Yes |
| **Number of paired volumes** | 40 | 40 | 50 | 40 | 1 |
| **Repetition Time (s)** | 4 | 4 | 3.8 | 3.5 | 4.886 |
| **Echo time (s)** | 0.01003 | 0.0029 | 0.017 | 0.0029 | 0.01 |
| **Post labelling delay (s)** | 1.8 | 1.25 | 1.517 | 1.5 | 2.025 |
| **Labeling duration(s)** | 1.8 | 1.5088 | 1.05 | 1.65 | 1.45 |
| **Dimension** | 64 x64 x32 | 96x96x40 | 64x64x24 | 64x57x16 | 128x128x40 |
| **Resolution (mm)** | 3.75x3.75x3.75 | 2.29x2.29x6 | 3.4x3.4x5 | 4x3.9x6 | 1.875x1.875x4 |
| **Magnetic Strength (T)** | 3 | 3 | 3 | 3 | 3 |
| **Flip angle ($^0$)** | 90 | 90 | 90 | 90 | 111 |

### Summary

- Subject ID: 99949
- Structural images: 1 T1-weighted
- ASL series: 1
    - Task: rest (1 run)
- Standard output spaces: MNI152NLin2009cAsym
- Non-standard output spaces:

### Arterial Spin Labelling

#### Reports for: task rest.

##### Summary

- Repetition time (TR): 4.3s
- Phase-encoding (PE) direction: Anterior-Posterior
- Slice timing correction: Not applied
- Susceptibility distortion correction: None
- Registration: FSL `flirt` rigid registration - 6 dof
- Confounds collected: std_dvars, dvars, framewise_displacement, trans_x, trans_y, trans_z, rot_x, rot_y, rot_z
- Motion summary measures: FD : 0.2214, relRMS: 0.001
- Coregistration quality: Dice Index: 0.9811, Jaccard Index: 0.963, Cross Cor.: 0.9771, Coverage: 1.0
- Normalization quality: Dice Index: 0.9602, Jaccard Index: 0.9234, Cross Cor.: 0.9499, Coverage: 0.9609
- Quality evaluation index : cbf: 0.8239,score: 0.8239,scrub: 0.8749, basil: 0.8647, pvc: 0.8086
- Mean CBF (mL 100/g/min) : GM CBF: 49.76, WM CBF: 39.15, GM/WM CBF ratio: 1.27
- Percentage of negative voxel : cbf: 0.14, score: 0.15, scrub: 0.05, basil: 0.0, pvc: 0.0

##### Alignment of asl and anatomical MRI data (surface driven)

FSL `flirt` was used to generate transformations from EPI-space to T1w-space - The white matter mask calculated with FSL `fast` (brain tissue segmentation) was used for BBR. Note that Nearest Neighbor interpolation is used in the reportlets in order to highlight potential spin-history and other artifacts, whereas final images are resampled using Lanczos interpolation.

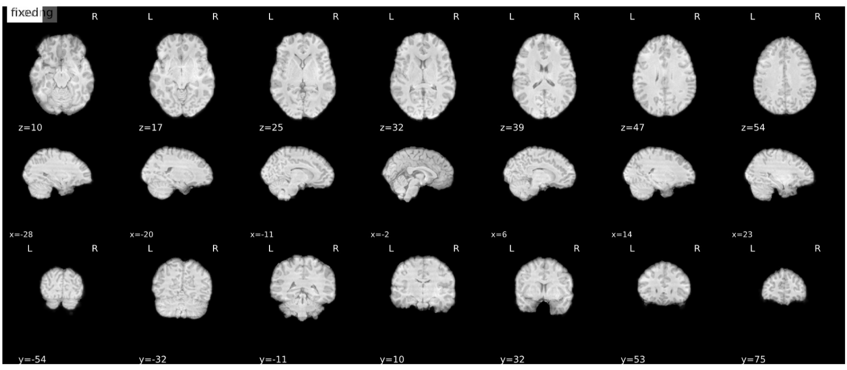


##### ASL Summary

Summary statistics are plotted, which may reveal trends or artifacts in the asl data. DVARS and FD show the standardized DVARS and framewise-displacement measures for each time point. A carpet plot shows the time series for all voxels within the brain mask. Voxels are grouped into cortical (blue), and subcortical (orange) gray matter, cerebellum (green) and white matter and CSF (red), indicated by the color map on the left-hand side.

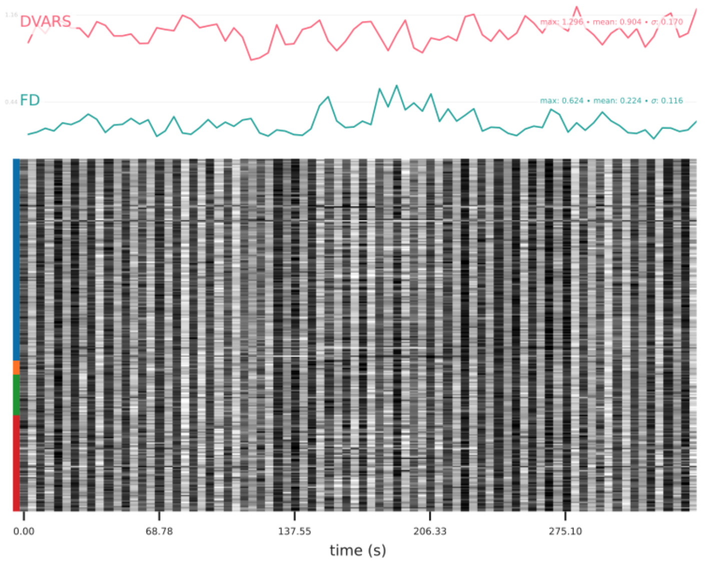


Get figure file: sub-99949/figures/sub-99949_task-rest_desc-carpetplot_asl.svg

##### CBF Summary

This carpet plot shows the time series for all voxels within the brain mask for CBF. Voxels are grouped into cortical (blue), and subcortical (orange) gray matter, cerebellum (green), white matter and CSF (red), indicated by the color map on the left-hand side. The score Index with value greater than zero indicates which volume(s) are removed by SCORE.

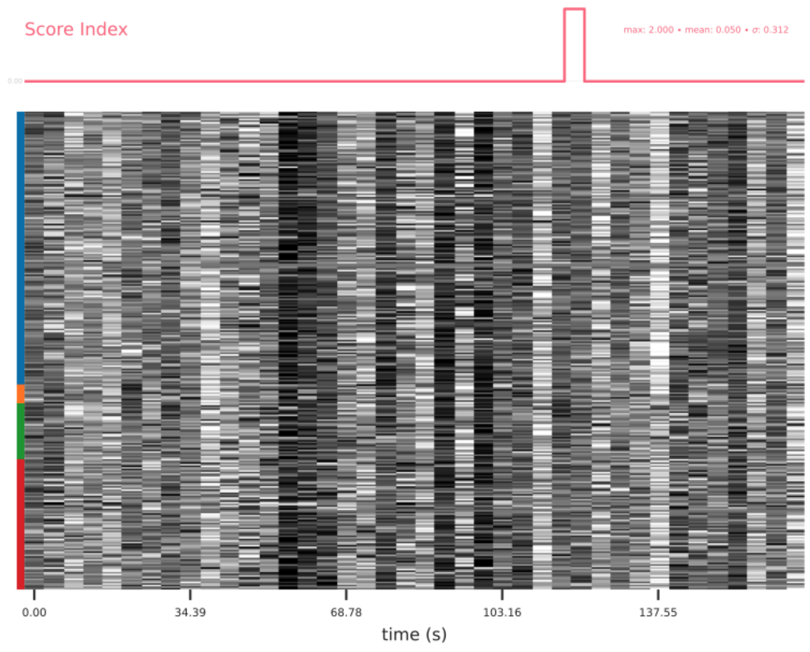


Get figure file: sub-99949/figures/sub-99949_task-rest_desc-cbftsplot_asl.svg

##### CBF maps

The maps plot cerebral blood flow (CBF) for basic CBF. The unit is mL 100/g/min

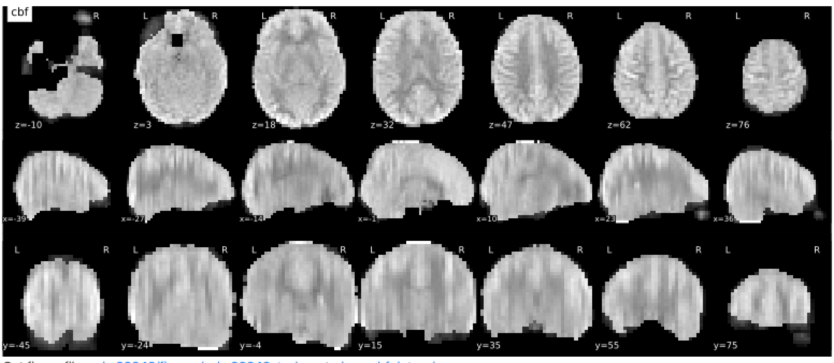


Get figure file: sub-99949/figures/sub-99949_task-rest_desc-cbfplot_asl.svg

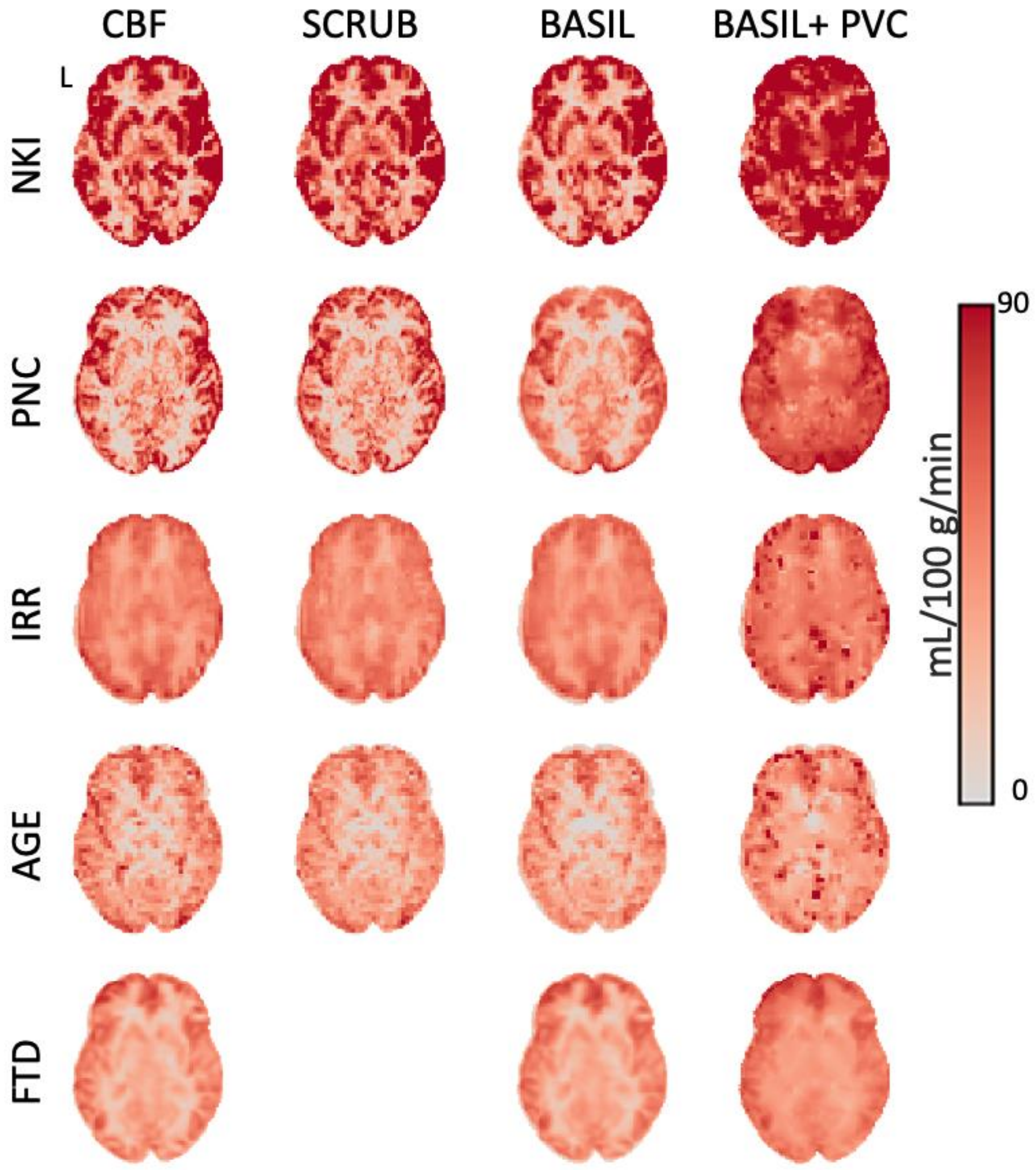


**Supplementary Figure 2 | Exemplar data for each dataset and CBF quantification method.** A single participant from each dataset is shown, with CBF quantified using each of four methods. SCRUB could not be applied for the FTD dataset as an ASL timeseries is required; the sequence used for that study provided only a ΔM image only.

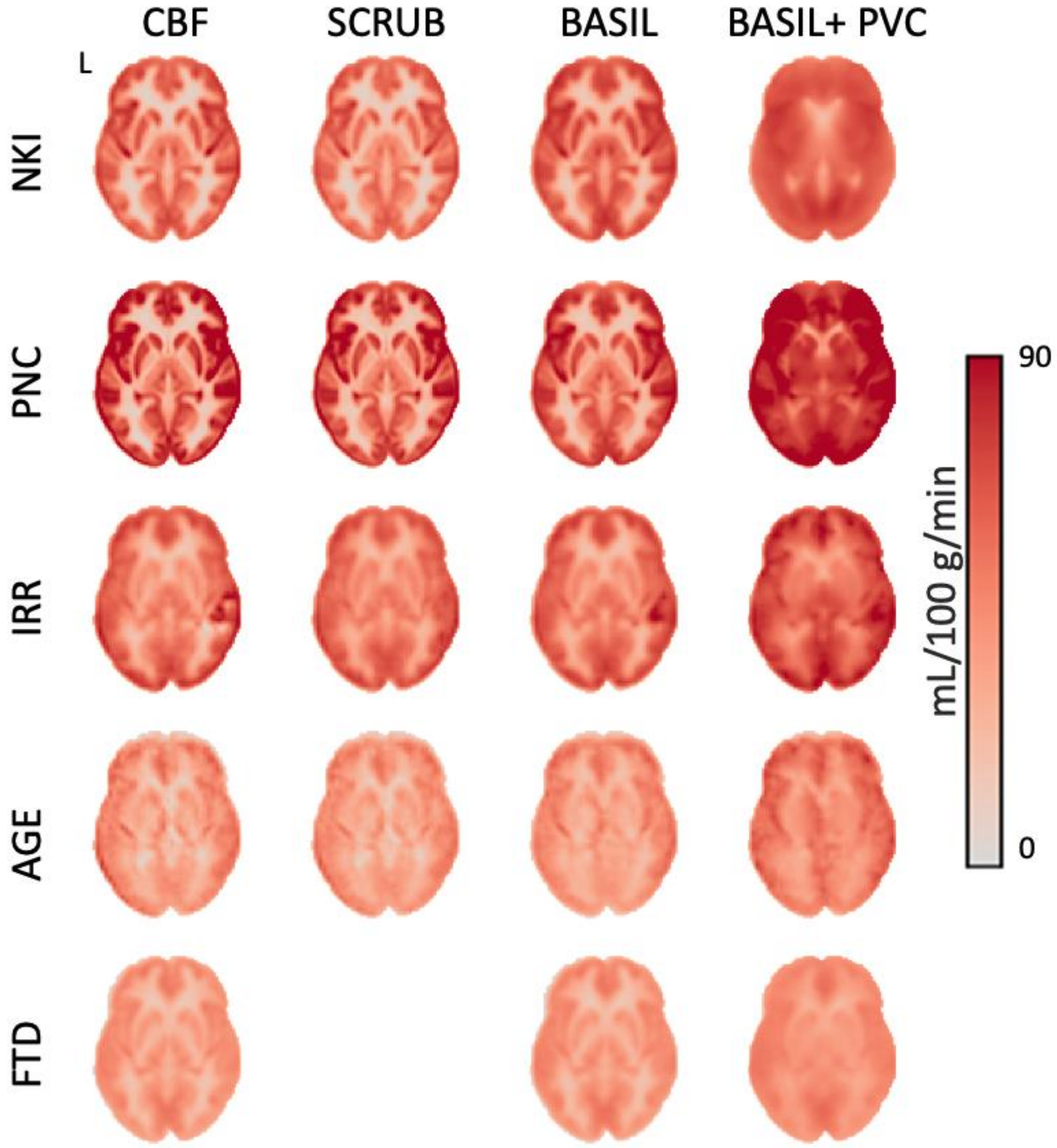


**Supplementary Figure 3 | Mean cerebral blood flow maps for each dataset and quantification method**. We quantified CBF for each dataset using four methods supported by ASLPrep. An axial slice (z=0) of the mean CBF image for each dataset is displayed for each quantification method. SCRUB could not be applied for the FTD dataset as an ASL timeseries is required; the sequence used for that study provided only a single ΔM image.

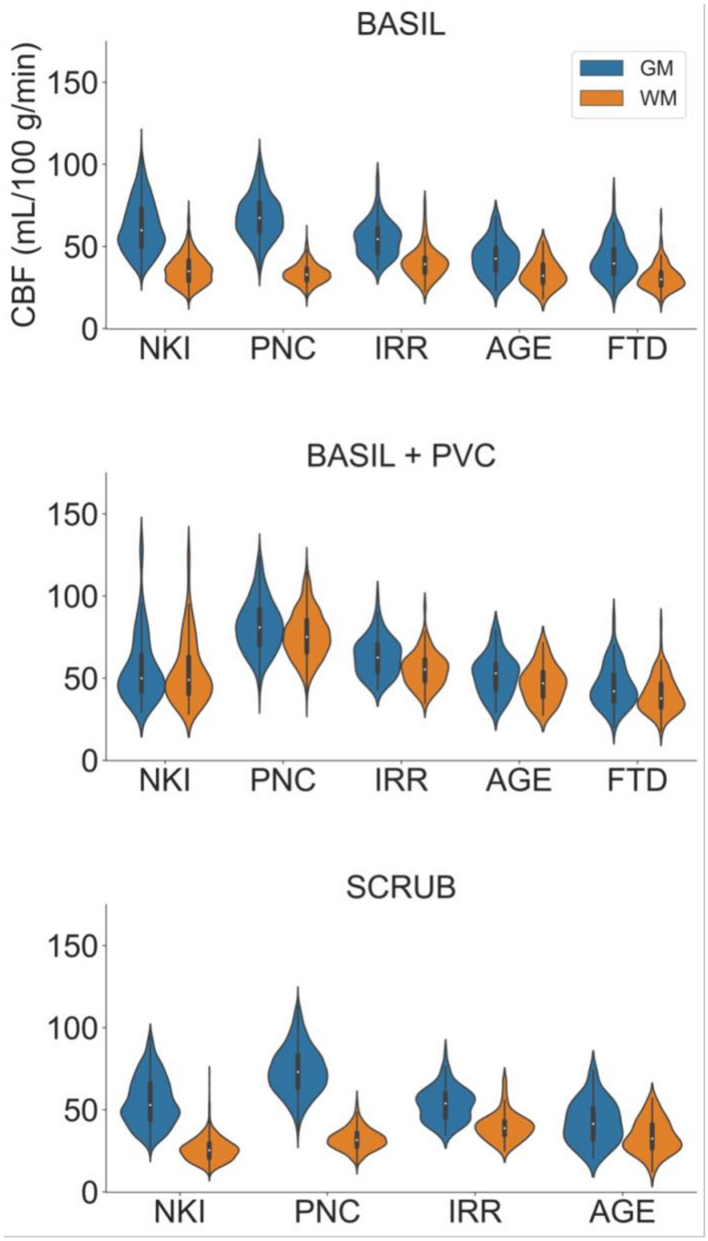


**Supplementary Figure 4 | CBF of gray and white matter across datasets.** The distribution of cerebral blood flow (CBF) within grey matter (GM) and white matter (WM) is displayed for each dataset, for each quantification option: the standard CBF model (see main text), BASIL, BASIL with partial volume correction (PVC), and SCRUB. SCRUB could not be applied for the FTD dataset as an ASL timeseries is required; the sequence used for that study provided only a single ΔM image

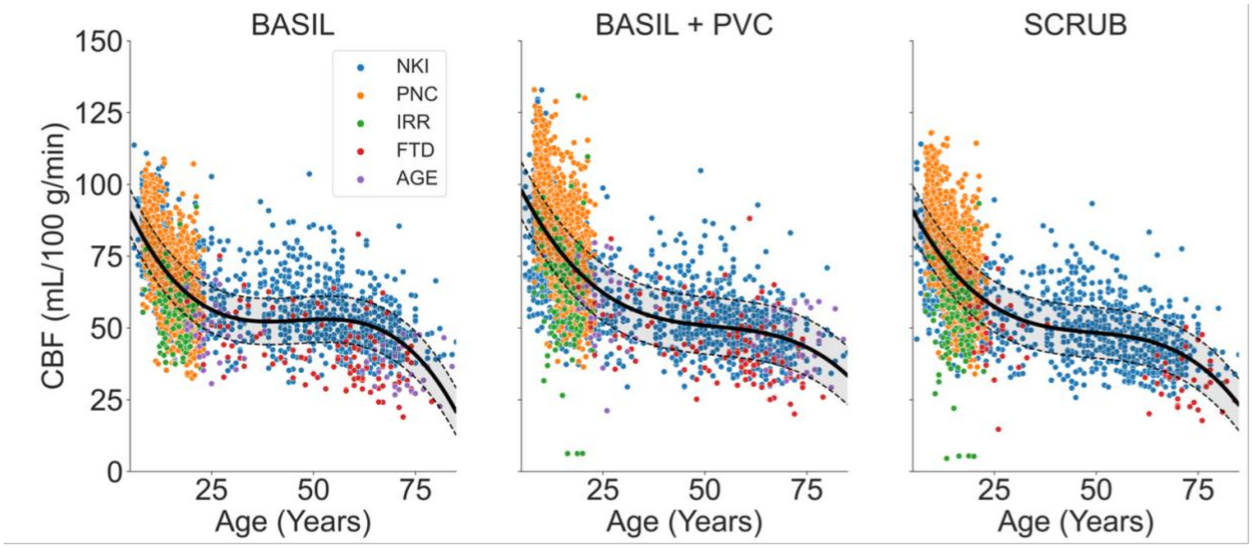


**Supplementary Figure 5 | CBF declines nonlinearly with age over the lifespan.** We evaluated how mean gray matter CBF evolved with age over the lifespan across all five datasets. For each dataset, we used four methods for quantifying CBF: the standard CBF model (see main text), BASIL, BASIL with partial volume correction (PVC), and SCRUB. We used a generalized additive model with penalized splines to characterize the nonlinear evolution of CBF over age. As expected, CBF declined rapidly in childhood, was fairly stable in adulthood, and declined again after age 60. Note that PVC results in increased CBF values, as expected.

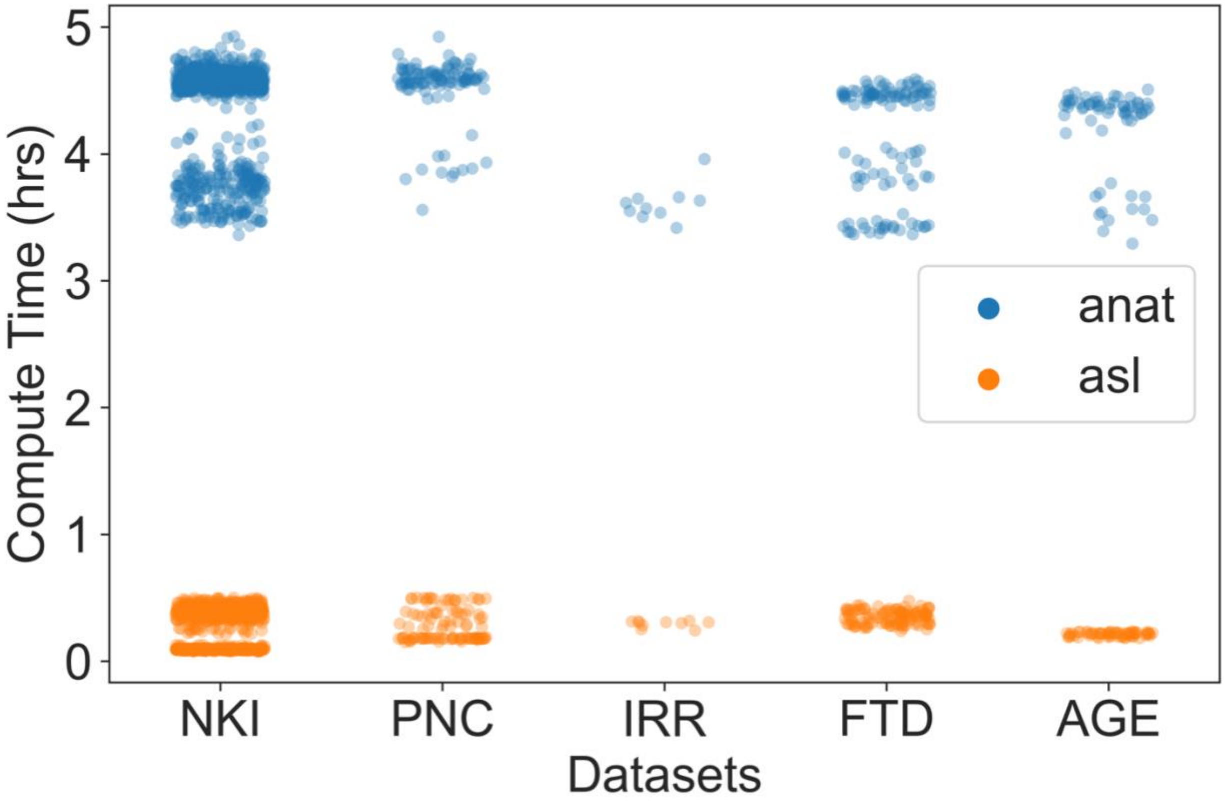


**Supplementary Figure 6 | Compute time for ASLPrep.** Distribution of compute time for each dataset, separated by ASL preprocessing and anatomic processing (which relies upon sMRIPrep). Anatomic preprocessing always required a longer duration, with ASL preprocessing and CBF computation requiring less than 40 minutes in all datasets.
